# 50 Hz volumetric functional imaging with continuously adjustable depth of focus

**DOI:** 10.1101/240069

**Authors:** Rongwen Lu, Masashi Tanimoto, Minoru Koyama, Na Ji

## Abstract

Understanding how neural circuits control behavior requires monitoring a large population of neurons with high spatial resolution and volume rate. Here we report an axicon-based Bessel beam module with continuously adjustable depth of focus (CADoF), which turns frame rate into volume rate by extending the excitation focus in axial direction while maintaining high lateral resolutions. Cost-effective and compact, this CADoF Bessel module can be easily integrated into existing two-photon fluorescence microscopes. Simply translating one of the relay lenses along its optical axis enabled continuous adjustment of the focal length. We used this module to simultaneously monitor activity of spinal projection neurons extending over 60 μm depth in larval zebrafish at 50 Hz volume rate with adjustable imaging thickness.

## 1. Introduction

Understanding how neural networks integrate inputs and generate outputs is a central goal of neuroscience. Neurons within these networks are distributed in three dimensions (3D). Monitoring their activity dynamics thus requires an imaging modality capable of rapid volumetric rates. Two-photon laser-scanning microscopy (2PLSM), combined with calcium indicators, has become the gold standard tool for *in vivo* monitoring of neural activity [1-3]. Calcium indicators with fast dynamics (rising time of ~40 ms for the dye Cal-520 [4, 5] and 80 ms for GCaMP6f [6]) can be used to monitor the firing of individual neurons to understand information flow in a network. However, capturing calcium dynamics of all labeled neurons in a volume within tens of milliseconds is challenging for 2PLSM because it images a volume by serially scanning the excitation focus in 3D.

Recently, we demonstrated a Bessel beam module that turns the frame rate of 2PLSM into volume rate by elongating its excitation focus in the axial direction [7]. The elongated, Bessel-like focus allowed us to image calcium dynamics of dendritic spines in 3D *in vivo* at 30 Hz, an unprecedented combination of high lateral resolution and fast volume rate. In this previous work, we employed a spatial light modulator (SLM) to generate the Bessel focus. Although such an SLM-based Bessel module can flexibly generate Bessel foci with variable lengths, numeric apertures (NA), and engineered axial profiles, it can be a challenge to fit a bulky SLM into existing microscopes with tight space. In contrast, an axicon (conical lens)-based Bessel module, a common method to generate an axially elongated focus, is less expensive and more compact [8-13], but can only generate a Bessel focus of a fixed axial length. In real-life neurobiological inquiries, however, it is often desirable to use Bessel foci of different axial lengths to investigate brain volumes of various sizes and/or labeling densities. To generate a Bessel focus with a different length, published axicon-based implementations modified the beam size, employed another axicon, or built an alternative optical path, all of which complicate the design and sacrifice the compactness of the axicon-based system [10, 11].

To address these limitations, here we introduce an axicon-based Bessel module with Continuously Adjustable Depth of Focus (CADoF). Simply by translating a lens within this module along its optical axis, we can continuously adjust the axial length of the resulting Bessel focus. We characterized the performance of this flexible module and demonstrated its usability by imaging groups of spinal projection neurons in the midbrain and hindbrain of zebrafish larvae [14-18], and achieved 50 Hz simultaneous volumetric activity imaging of these neurons during escape behavior.

## 2. Materials and Methods

### 2.1 Construction of the axicon-based CADoF Bessel module

The axicon-based CADoF module was built between a Ti-Sapphire laser (Mai Tai HP, Spectra-Physics) and a custom two-photon fluorescence microscope equipped with a resonant galvanometer and controlled by ScanImage2016 (Vidrio Technologies, LLC). The axicon was housed in a lens tube (SM1L05, Thorlabs), which was in turn mounted in a translational cage (CXY1, Thorlabs). The axicon was placed at the front focal plane of a lens (L1 in **Fig. 1a**) with 80-mm focal length (AC254-080-B-ML, Thorlabs), which transformed the light refracted by the axicon into a ring. An annular aperture mask was placed at the back focal plane of the same lens to spatially filter the ring. A lens pair (L2 and L3 in **Fig. 1a**, AC254-150-B-ML and AC254-200-B-ML, Thorlabs) relayed the ring onto the galvo scanners and then to the back focal plane of the objective. Both the axicon and the lenses were built within a caged system, allowing easy translation of lens L2 along its optical axis. Two mirrors mounted on rail assemblies (16.011.0130, 16.021.0065 and 16.021.0020, OWIS, Germany) were used to switch between Bessel and Gaussian focus scanning. During system characterization, three convex axicons with apex angle of 178° were tested. Axicon 1 and Axicon 2 were from the same vendor (1-APX-2-H254-P, ALTECHNA Inc.). Axicon 3 was from ASPHERICON (XFL25-010-U-B).

**Figure 1.**
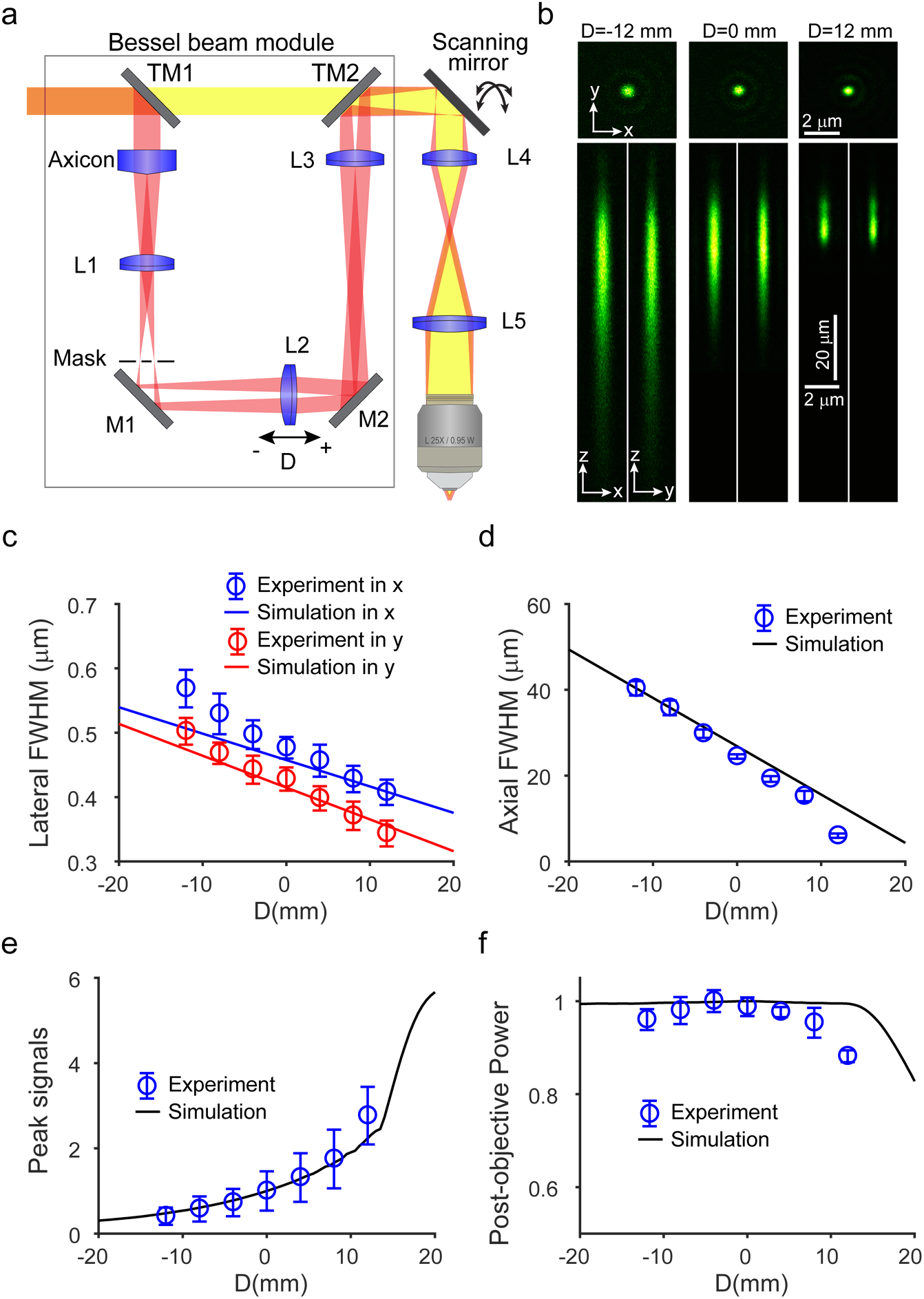
Design and characterization of the axicon-based CADoF Bessel module. **(a)** Optical schematics of the Bessel module incorporating an axicon. Two translating mirrors (TM1 and TM2) switch between the Gaussian and Bessel imaging modalities. D is defined as 0 mm when the mask is at the front focal plane of L2, and positive when L2 moves away from the mask. **(b)** Experimental axial PSFs with D = -12 mm, 0 mm, and 12 mm, respectively. Varying D leads to variable **(c)** lateral FWHM, **(d)** axial FWHM, **(e)** peak signal of PSF, and **(f)** post-objective power. Data in (**c-e**) were collected from nine 0.2-μm-diameter beads, and data in **(f)** are averages of four measurements. Lateral FWHMs in **(c)** were measured at the peak axial locations. Error bars represent standard deviations. Simulation data in **(c)** and **(d)** were linearly fitted (R^2^ > 0.995). Excitation light was linearly polarized along x axis for simulation. Focal lengths of L1 to L5: 80 mm, 150 mm, 200 mm, 35.2 mm, and 200 mm, respectively. The apex angle ofthe axicon was 178°. The 1/*e*^2^ diameter ofthe beam at the axicon was 2.8 mm. Objective: Leica, 0.95 NA, 25X; wavelength: 940 nm. The annular aperture mask had an outer diameter of 1.312 mm and an inner diameter of 1.208 mm.

### 2.2 Design of the annular aperture mask

Annular aperture masks with different annular thicknesses and diameters were designed. **Supplementary Code** “maskDesign.m” (Matlab) generates four masks with the inner and the outer diameters located at 1/e, 1/(5e), 1/(10e) and 1/(15e) of the peak amplitude of the ring, respectively. The transmittance ratios of these four masks under ideal conditions are 92.4%, 99.0%, 99.6% and 99.8%, respectively (see **Supplementary Code** “demo1.m”).

### 2.3 Zebrafish preparation

All animal experiments were conducted according to the National Institutes of Health guidelines for animal research. Procedures and protocols on zebrafish were approved by the Institutional Animal Care and Use Committee at Janelia Research Campus, Howard Hughes Medical Institute. Larval zebrafish *(Danio rerio)* of pigment mutant *casper* [19], which is a compound mutant of *mitfa*^w2/w2^ *(nacre)* and *roy* ^a9/a9^, was raised at 28.5 °C with a standard procedure [20]. At 5 days post fertilization, 25% w/v solution of Cal520-Dextran conjugate (10,000MW, AAT Bioquest, California) in zebrafish extracellular solution (134 mM NaCl, 2.9 mM KCl, 1.2 mM MgCl_2_, 2.1 mM CaCl_2_, 10 mM HEPES, and 10 mM glucose, adjusted to pH 7.8 with NaOH) was pressure injected into the ventral spinal cord at the level of 20^th^ myotome segment following standard procedures [21]. The larvae were left in filtered fish system water at 28.5°C overnight to allow retrograde filling of spinal projection neurons (reticulospinal and vestibulospinal neurons). On the next day, before imaging, larvae were briefly anesthetized by bath application of 0.02% w/v solution of Ethyl 3-aminobenzoate methanesulfonate (Sigma-Aldrich, St. Louis) in filtered fish system water for 1 min. Larvae were mounted dorsal-side up in a small drop of 1.6% low melting point agarose (Invitrogen) on a glass-bottomed dish. The agar covering the caudal body was removed so that the trunk movement could be recorded. Anesthetic was replaced by filtered fish system water. During imaging, the dish containing the fish was tightly glued on the top of a stimulus platform with dental wax (GC Corporation). A loudspeaker (FRS-8, Visaton) was mounted on the platform. Sinusoidal stimulus waveforms (500 Hz, 2 cycles) were generated by a function generator (AFG3102C, Tectronix), amplified by an audio amplifier (LP-2020A, Lepy), and delivered to the loudspeaker. Images were acquired from 2 s before to 5 s after the stimulus with inter-stimulus interval of longer than 1 min. Blood flow and tissue transparency were inspected after the imaging experiments to confirm that two-photon excitation beam did not damage the brain tissue.

### 2.4 Data analysis

All functional images were registered using an iterative cross-correlation-based registration algorithm [22]. ROIs were manually outlined and then compared with Gaussian 3D image stack for the anatomy. For the calculation of the calcium transient ΔF/F, each ROI across different trials shared the same baseline, determined as the mean calcium fluorescence signal in the 1-sec window prior to the stimulus. Those trials that had spontaneous calcium transients within this pre-stimulus window were excluded from baseline calculation. A neuron was defined as responsive if the amplitude of the maximal calcium transient was bigger than the mean + 3SD of the baseline. Scan time lag resulting from different cell positions and the vertical scanning of y galvo was corrected according to vertical positions of centroids of individual ROIs.

## 3. Results

### 3.1 Axicon-based CADoF module

To generate continuously adjustable Bessel foci, we started with the optical path similar to conventional axicon-based Bessel beam module [10, 11]. Placed between the light source and the microscope (**Fig. 1a**), the key component of the module is an axicon. The axicon has a conical surface, where light is refracted according to Snell’s law. The lens L1 then shapes the light into a ring at its back focal plane, which is in turn projected by a lens pair consisting of L2 and L3 to the galvo scanning mirrors, and then to the back focal plane of the objective by another lens pair L4 and L5. The annular aperture mask placed at the back focal plane of L1 is optional, but helps to shape the axial profile of the two-photon excitation point spread function (PSF) by blocking the unwanted light that arises from the imperfections of the axicon. This optical design is conceptually similar to the traditional axicon-based modules [10, 11, 23]. The novelty of the present study, however, is that by moving the lens L2 along its optical axis, we can continuously vary the axial length of the Bessel focus (**Fig. 1** and **Visualization 1**). (Moving lens L3 also varies the axial length – for simplicity, we only discuss the effects of translating L2.) **Fig. 1b** shows three representative (long, medium, and short) axial PSFs with full-widths-at-half-maxima (FWHMs) of 39 μm, 24 μm, and 14 μm at lens L2 displacements of D=-12 mm, 0 mm, and 12 mm, respectively. When the displacement of L2 is negative (i.e., moving L2 closer to the mask), more power is allocated to the central region of the objective back focal plane (**Fig. 2**, 1st column), which yields a smaller effective NA and, together with the phase distribution of the pupil function, a longer Bessel focus. In contrast, moving the lens away from the mask distributes more power to the edge of the objective pupil function, leading to a bigger effective NA and a shorter focus (**Fig. 2**, 3rd column). This way, moving the lens from left to right in **Fig. 1a** allows us to continuously change the lateral and axial FWHMs (**Figs. 1c, d**). Numeric simulations using the vector diffraction theory developed by Richards and Wolf [24] agree well with the experimental data (**Figs. 1c-f**), where the difference between the lateral resolutions along x and y axes is caused by the linear polarization of the excitation light (**Fig. 1c**). It is worth noting that there is a tradeoff between axial/lateral FWHM and the peak signal. Given the same power at the axicon, longer Bessel foci (negative D values in **Fig. 1d**) have larger lateral FWHMs (**Fig. 1c**) and smaller peak signals (**Fig. 1e**), because there is less energy in the central peak of the foci. For the shortest Bessel foci, light starts to be clipped by the objective, leading to a decrease in the post-objective power (**Fig. 1f**).

**Figure 2.**
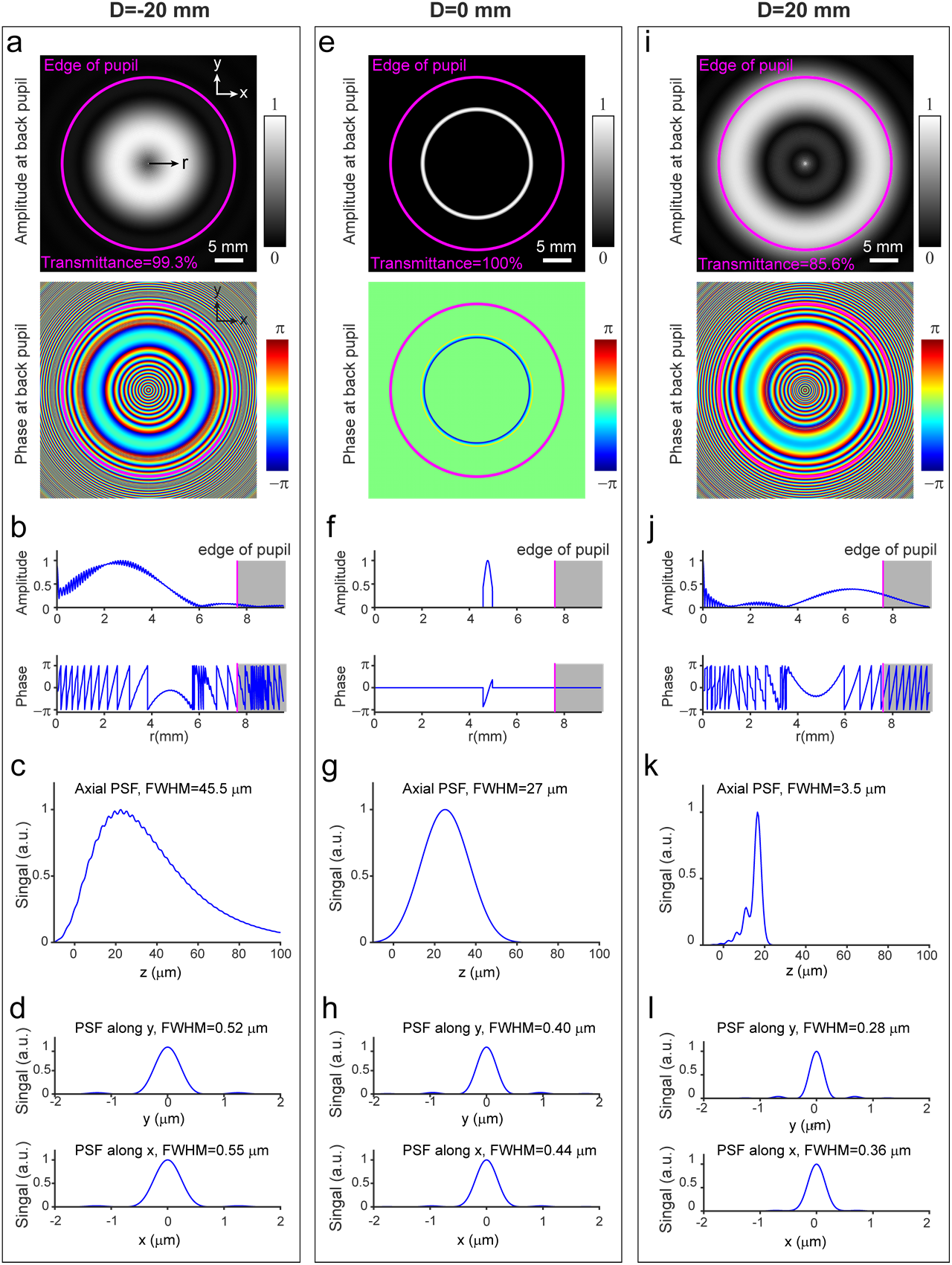
Engineering Bessel foci by displacing Lens L2. **(a)** 2D and **(b)** 1D (along x direction) representations of the amplitude and phase of the pupil function at the objective back focal plane, **(c)** axial and **(d)** lateral two-photon excitation PSFs for D = -20 mm, 0 mm, and 20 mm, respectively. Same parameters as described in the caption of **Fig. 1**. (See **Visualization 1** for pupil function and PSF during continuously displacing Lens L2 from - 20 mm to 20 mm.)

Good agreement between experimental data and theoretical predictions as shown in **Fig. 1** was achieved by having a thin annular aperture mask (e.g., 52 μm thickness used for data in **Fig. 1**) to correct for the imperfections of the axicon (**Fig. 3**) [13]. However, such a thin mask causes power loss (**Fig. 3**), which might become problematic in systems with limited excitation power. One can use a thicker mask or even operate without mask and obtain non-theoretical yet still desirable axial profiles. Another method to alleviate the deviation from theoretical predictions is to substantially expand the beam before the axicon [25, 26]. To maintain the compactness of the module, we did not adopt this strategy.

**Figure 3.**
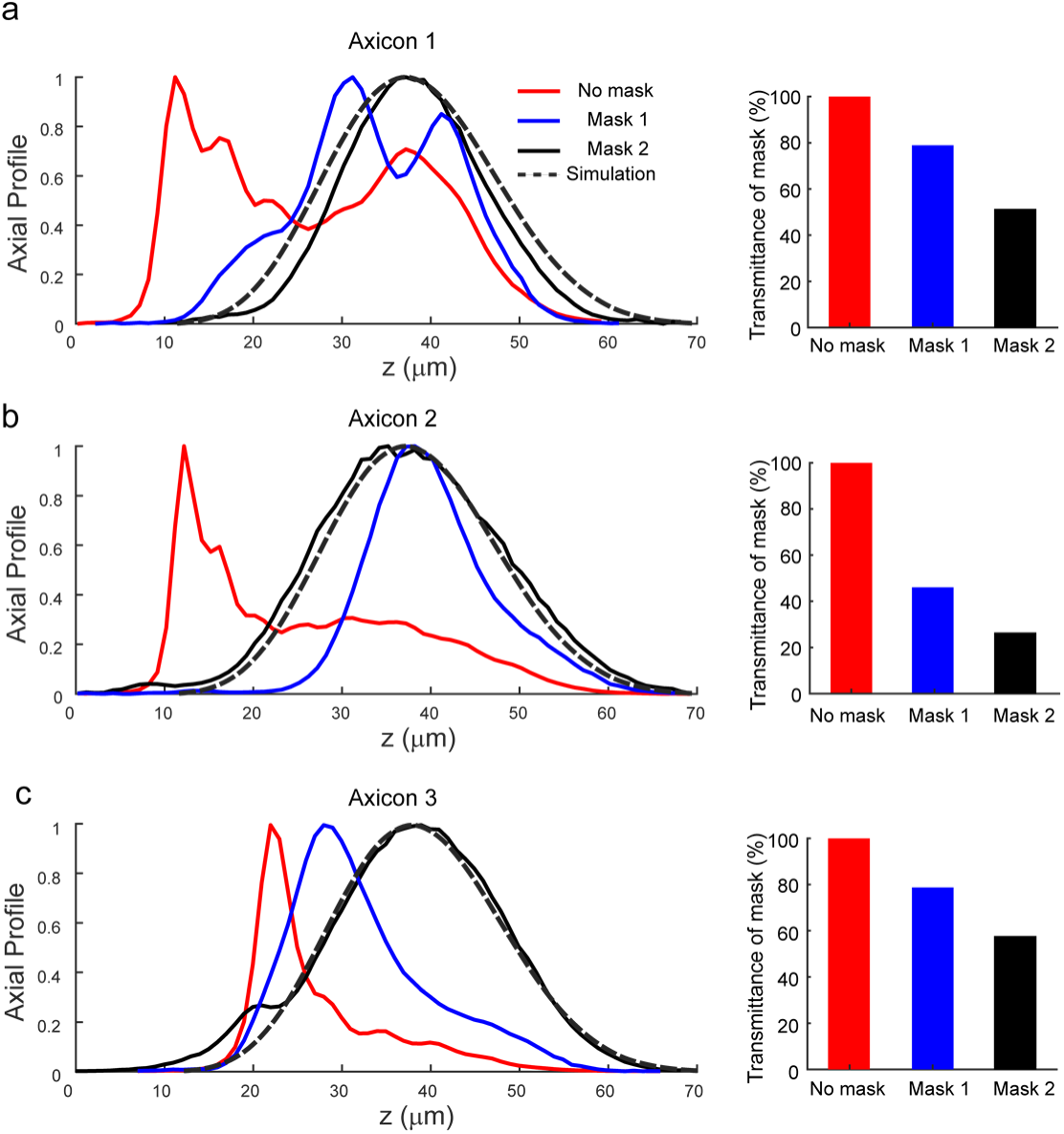
Annular aperture masks alter the axial profile of the Bessel focus. (a-d) the axial PSFs measured without mask (red line), with Mask 1 (blue line), with Mask 2 (black line), together with the simulated PSF (dashed line) for Axicons 1-4, respectively. All axicons have the same apex angle (178°). Axicons 1 and 2: 1-APX-2-H254-P, ALTECHNA; Axicon 3: XFL25-010-U-B, ASPHERICON. Outer diameter: 1.429 mm, inner diameter: 1.091 mm for Mask 1; Outer diameter: 1.312 mm, inner diameter: 1.208 mm for Mask 2. Theoretical transmission ratio through Mask 1 and Mask 2 for perfect axicons: 99% and 90%, respectively.

### 3.2 Axicon-based CADoF module enables flexible control over interrogated volume thickness in zebrafish larvae

To demonstrate the flexibility of the CADoF module, we first imaged groups of spinal projection neurons (reticulospinal and vestibulospinal neurons) in the midbrain and hindbrain of larval zebrafish. These neurons form a stereotypic organization arranged in a volume around 400 μm × 200 μm × 100 μm (rostrocaudal, mediolateral, and dorsoventral axes, respectively) and were retrogradely labeled with fluorescent tracer dye injected into the spinal cord, permitting easy identification of individual neurons from one zebrafish to another [18, 27]. Here we defined the origin of relative depth (z = 0 μm, 96 μm from the skin surface) as the plane where the first labeled structure appeared when the objective was moved from the dorsal to ventral part of the zebrafish. Scanning a Gaussian focus (axial FWHM: 2.6 μm) in 2D, we could only image a subset of neurons (e.g., at maximum, 38 neurons, **Fig. 4a**). To sample the 74-^m-thick volume containing all labeled neurons, 28 slices of 2D images have to be sequentially acquired (**Fig. 4b**). With the CADoF modality, even the shortest Bessel focus revealed more neurons without moving objective (arrowheads in **Fig. 4c**). As the Bessel focus continued to extend axially, more cells showed up successively (arrowheads in **Figs. 4d-j**). The longest Bessel focus revealed 108 neurons (**Fig. 4j**), with a lateral resolution that provided comparable cellular-resolving power to the mean-intensity projection image of the 2D Gaussian image stack along the dorsoventral axis (**Fig. 4b**).

**Figure 4.**
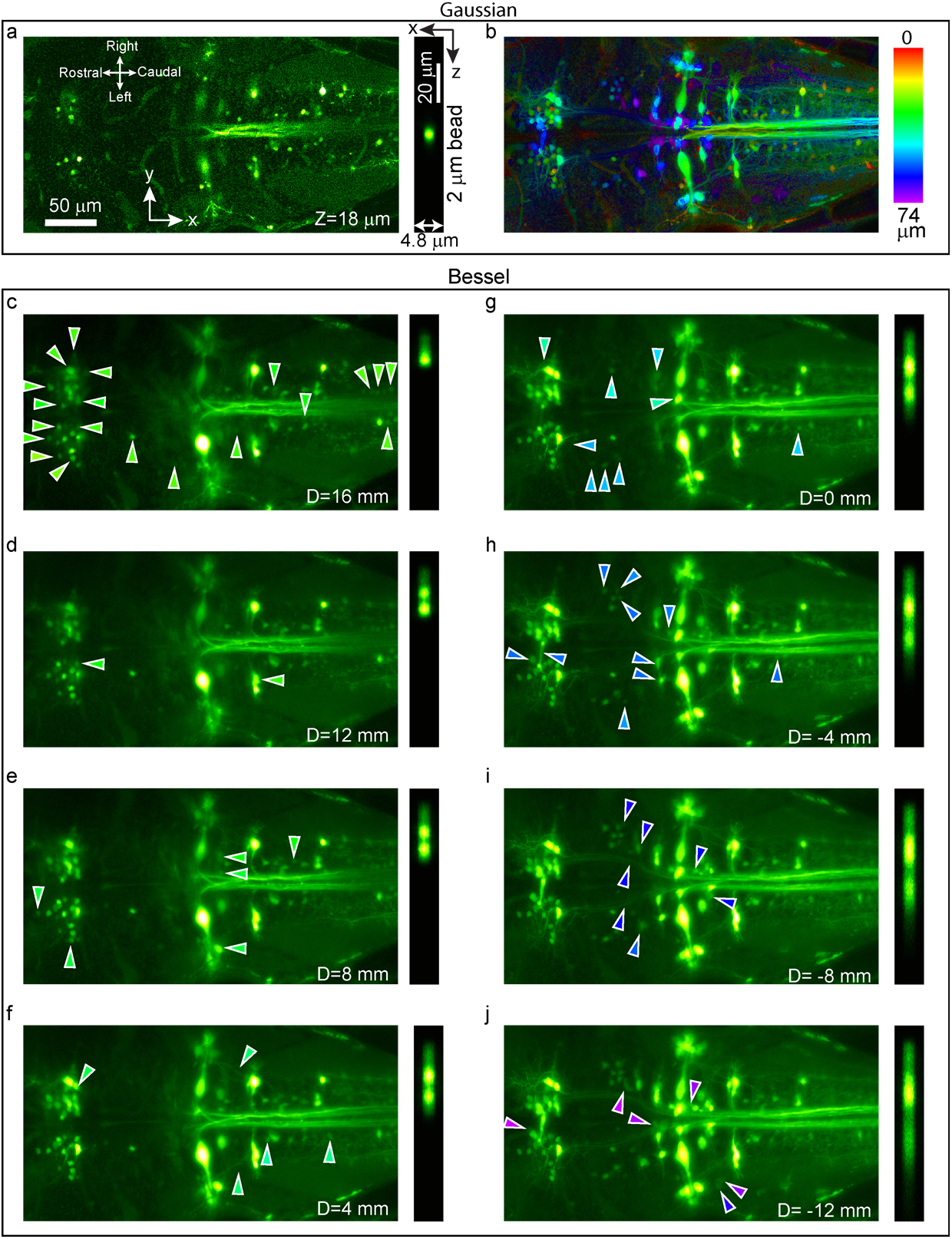
Scanning Bessel foci with variable axial lengths probes varying volumes of spinal projection neurons *in vivo*. Reticulospinal and vestibulospinal neurons labeled with Alexa Fluor 488 in a larval zebrafish were imaged by scanning either **(a,b)** Gaussian or **(c-j)** Bessel foci. Left panels: neuron images; Right panels: axial profiles of 2^m-diameter beads. **(a)** An image acquired using Gaussian focus at the relative depth of 18 μm (114 μm from the surface) contains 38 neurons. **(b)** Mean intensity projection over 74 μm axial range (absolute depth from 96 μm to 170 μm), color-coded by relative depths. **(c-j)** Volumetric images acquired by scanning Bessel foci with different axial lengths. Longer foci revealed more structures, e.g., **(c)** 65 neurons; **(g)**: 87 neurons and **(j)**: 108 neurons. Arrowheads (same color code as in b) in each image point to example new structures (neurons or axons) compared to the previous image. Axicon 1 and Mask 1 were employed here.

### 3.3 50 Hz volumetric calcium imaging of spinal projection neurons in zebrafish larvae

After demonstrating the performance and flexibility of our axicon-based CADoF Bessel module on structural imaging of the spinal projection neurons, we moved on to investigate the activity dynamics of these neurons in zebrafish larvae engaging in escape behaviors [14-17]. We retrogradely labeled these neurons with a calcium indicator dye (Cal-520 dextran) from the ventral spinal cord. Acoustomechanical tapping stimuli were applied during imaging to trigger escape response and neural activity (**Visualization 2**). In one zebrafish (**Fig. 5**), the labeled neurons were distributed in a volume of 366 μm × 214 μm × 66 μm (**Visualization 3**). Gaussian focus scanning over a single plane (**Fig. 5a**) revealed four active neurons (regions of interest, ROIs 1-4, **Fig. 5b**) including two Mauthner cells (ROIs 1 and 2, the largest bilateral pair of reticulospinal neurons known to initiate escape behaviors [28-30]) as reported in a previous study [31]. Switching to the CADoF module used in **Fig. 4**, we measured calcium activity from neurons distributed in volumes of varying thickness at 50 Hz volumetric rate. The short Bessel focus (14 μm axial FWHM, D = 12 mm, **Fig. 5c**) revealed the same four active neurons (**Fig. 5d**) with similar responses to those observed with Gaussian focus scanning. The medium Bessel focus (24 μm axial FWHM, D = 0 mm) uncovered six additional responsive neurons (ROIs 5-10 in **Figs. 5e,f**). Scanning the long Bessel focus (39 μm axial FWHM, D = -12 mm) covered the entire 366 μm × 214 μm × 66 μm volume and further revealed another five active neurons (ROIs 11-15 in **Figs. 5g,h**). In addition, the calcium transient signal-to-noises of several neurons increases by having a longer Bessel focus integrate the signal throughout their cell bodies (ROIs 3,5 and 9). Overall, for the 36 neurons that were labeled by Cal520 dextran in this fish (**Figs. 5i,j**), we monitored their activity at 50 Hz and identified 15 neurons that exhibited tapping-evoked calcium transients.

**Figure 5.**
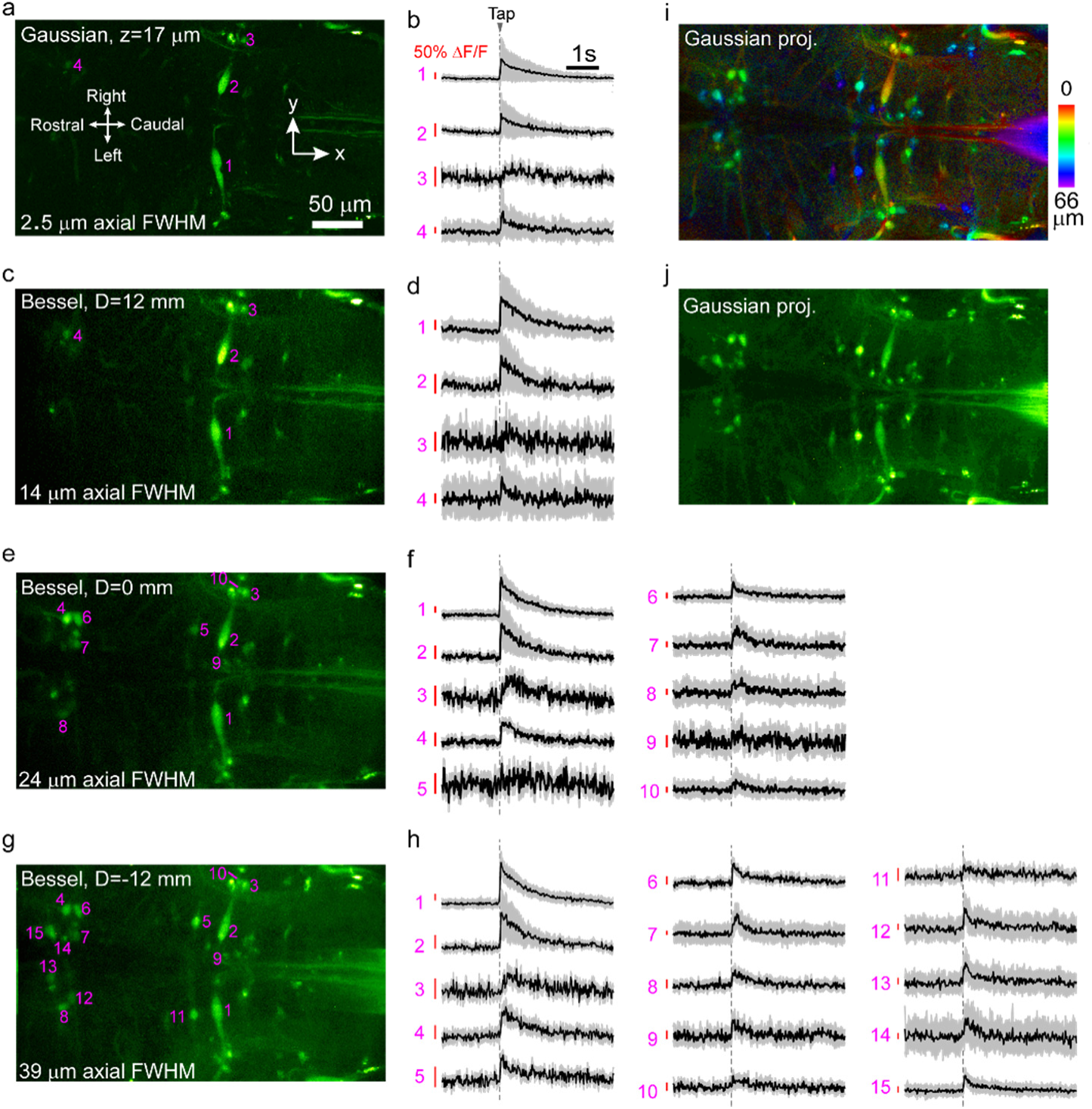
50 Hz volumetric functional calcium imaging of volumes of spinal projection neurons in zebrafish larvae. **(a)** Image acquired by Gaussian focus scanning at 127 μm from the dorsal surface of the head (relative depth z = 17 μm). **(b)** Averaged calcium transients of neurons evoked by the acoustomechanical tapping stimuli. **(c)**, **(e)**, and **(g)** were volumetric images obtained by scanning a short (14 μm axial FWHM), medium (24 μm axial FWHM), and long (39 μm axial FWHM) Bessel foci, respectively. **(d)**, **(f)**, and **(h)** were averaged calcium transients of responsive neurons. An acoustomechanical stimulus was delivered at 0 s. The Gaussian focus had 2.6 μm axial FWHM. The short (displacement of lens L2 D = 12 mm), medium (D = 0 mm) and long Bessel (D = -12 mm) foci has 14 μm, 24 μm and 39 μm axial FWHMs. The field of view was 366 μm × 214 μm. (See **Visualization 2** for the functional movies.) **(i)** and **(j)** Mean intensity projections of a 66-μm-thick image stack acquired by Gaussian focus scanning (see **Visualization 3** for the Gaussian 3D stack). Color in **(i)** encodes relative depth. Eleven trials were averaged in **(b)**, **(d)**, **(f)**, and **(h)**. Shadow represents standard deviations. Postobjective power: **(a)** 38 mW, **(c)** 97 mW, **(e)** 110 mW and **(g)** 132 mW.

## 4. Discussions

Bessel focus scanning for volumetric imaging provides several advantages: reduction of data size, resistance to artifacts induced by axial motions, compatibility with other volumetric imaging method [32], and easy integration into existing microscopes as a standalone module [7]. In practice, it is desirable to have the length of Bessel focus continuously adjustable: For samples with sparse labeling, a long (e.g., > 100 μm) Bessel focus can monitor activity over a larger volume. For more densely labelled samples, a shorter Bessel focus (e.g., 30 to 80 μm) can reduce the occurrences of overlapping structures. An even shorter (e.g., 20 μm) Bessel focus is sufficient when used to eliminate axial-motion-induced image artifacts during experiments with behaving animals. Both SLM-based [7] and axicon-based CADoF (**Fig. 1**) Bessel modules can generate Bessel foci of continuously tunable axial lengths. SLM-based Bessel beam module achieves this by varying the phase pattern loaded on the SLM [7]. Here, we showed that displacing the Lens L2 in **Fig. 1** in the axicon-based CADoF Bessel module changed the electric field distribution at the objective back focal plane (**Fig. 2**), where variations in both the amplitude and the phase of the electric field work concertedly to generate Bessel foci of different axial and lateral widths. (If we maintain the amplitude distribution but force the phase of the electrical field on the back pupil plane to be constant, the resulting pupil function fails to produce a Bessel focus, **Visualization 4**). Although the module described in this paper does not have the same flexibility in engineering the PSF profiles as SLM-based modules do [7, 33], this axicon-based Bessel module costs much less to set up (~$5,000, rather than $30,000, for the entire module), occupies less space (due to its transmissive layout), and is insensitive to polarization of the light, making it an attractive alternative to the more expensive modules.

We used the axicon-based CADoF Bessel module for 50 Hz volumetric calcium imaging of spinal projection neurons in zebrafish larvae during behavior (**Figs. 4, 5**). Because more energy is distributed in the side rings of the Bessel foci, to get the same signal, higher average power is required for Bessel foci than for Gaussian ones. We did not observe damages on brain tissue, even after dozens of imaging trials. This may be attributed to two factors: the amount of two-photon excitation, as reflected by the final fluorescence signal, stayed similar, indicating that Bessel focus scanning does not introduce additional photo-excitation-related damages; the absorption of the 940 nm light by the larval brain and the immersion media was minimal, thus leading to minimal heating-related damages.

The ability to measure the activity of all neurons within the image volume simultaneously using the axicon-based CADoF Bessel module, instead of plane by plane as done in conventional approaches, would make it possible to study the neural correlates of complex behavior repertoire in single trials. For sensors of neural activity [34] (e.g., for neurotransmitter release or membrane voltage) whose transients are much shorter in time than calcium sensors, the ability to image a volume at tens of Hz as we demonstrated here would be essential to capture all the activity events during behavior. For targeting neurons within smaller fields of view, one can image at even faster volume rate by reducing number of scanning lines. Beyond neural imaging, this axicon-based CADoF Bessel module can benefit other applications where an extended focus is used for optical measurement or manipulation.

## Acknowledgements

The authors thank Anderson Chen for suggestions on Bessel module construction, Yajie Liang for help with optical system alignment, Janelia Experiment Technology for designing and building of the microscope, and vivarium staff for fish care.

